# Selective Laser Etching technology for reconfigurable microfluidic and electrochemistry-on-chip

**DOI:** 10.64898/2026.01.08.698316

**Authors:** Inès Muguet, David Bourrier, Pierre-François Calmon, Pierre Lapèze, Pierre Joseph, Morgan Delarue

## Abstract

Polydimethylsiloxane (PDMS) is widely used in academic microfluidics due to its favorable biocompatible properties and compatibility with soft lithography. Moreover, the recent developments of reconfigurable microfluidics rely on the microfabrication of sliding elements, which are 3D objects insertable inside a microfluidic chip to provide a given function. However, the complexity of microfluidic device geometries or sliding elements remains largely limited by the traditional microfabrication methods such as, among others, on SU-8 photolithography or dry epoxy films. Such methods are suited for simple, planar “2.5D” structures with uniform depths, yet struggle to produce more advanced architectures due to material and alignment constraints. To address these limitations, we propose to investigate advanced laser-based approaches, such as Direct Laser Writing (DLW) and Selective Laser Etching (SLE), which enable the construction of high-resolution, 3D designs. We further explore the use of SLE to fabricate sliding elements that enhance chip functionality, including chamber reconfiguration and biological sample manipulation. As a proof of concept, we show that these elements can be functionalized into pH microsensors, paving the way to reusable and reconfigurable electrochemistry-on-chip.

## INTRODUCTION

In the world of academic microfluidics, silicone-based polymers, particularly polydimethylsiloxane (PDMS), have long been a go-to material for chip fabrication. Its popularity stems from a unique combination of physical and chemical characteristics: optical clarity across the visible spectrum, minimal autofluorescence, elasticity, permeability to gases, and biocompatibility^1–5^. The microfluidic chips fabrication process typically involves pouring liquid PDMS onto a mold, where it cures and conforms with high accuracy to fine features. This microreplication protocol, designed as soft lithography^6–8^, allows for transferring the micro-to nanoscale structures from the mold^9^, thanks to the flexibility of the PDMS and its minimal shrinkage. Once cured, the polymer can be bonded to glass, silicon, or additional PDMS layers, often using plasma treatment, to construct sealed microchannels.

Traditional mold fabrication often depends on conventional or multi-level photolithography techniques and commonly used materials include SU-8 patterned substrates, dry films or various thermoplastics^10^. Such additive fabrication methods are well-suited for creating straightforward designs, such as linear channels, basic geometric features and uniform channel depth, but are not suitable for more complex microchannels geometries, typically due to alignment difficulties. Consequently, the microchannels produced using these techniques are not really 3D and characterized by features like straight-edge, rectangular profiles and uniform heights. It is possible to achieve slightly more complex shapes, such as hemispherical channels or varying channel depths^11,12^, yet these “quasi-3D” geometries are typically limited and require additional steps when using conventional fabrication approaches.

Moreover, the advent of reconfigurable microfluidic rely on the use of sliding elements to bring novel functions inside a PDMS chip^1–4^. Such elements are microfabricated rods that can be inserted inside microfluidic systems, enabling a variety of applications: creation of tailored and reconfigurable culture chambers, biological sample loading and retrieval, biological sample placement and aspiration, creation of pressurized culture chambers. Currently developed sliding elements are usually produced using epoxy resist-based materials, which have the disadvantage to be auto-fluorescent. As a result, their fabrication process encounters similar issues to conventional photolithography ones, and the structures generally remain quasi-3D to achieve high resolution^2^.

To overcome such limitations, novel methods are being investigated for the fabrication of complex microfluidic molds^10^. For instance, Selective Laser Etching (SLE) techniques use femtosecond laser irradiations to induce local modifications in a transparent material, typically fused silica substrates (amorphous SiO_2_). These local modifications make the material more sensitive to chemical etching. Thus, subsequent chemical treatment etches out laser modified areas to reveal the desired pattern.

Such components are becoming increasingly important in fields like sensors, microfluidics, medical technology, data storage, and photonics^13–15^. Their popularity stems from the silica exceptional thermal, chemical, and mechanical resistance, combined with low thermal expansion and excellent optical properties. These microstructures typically range in size from a few millimeters down to a few micrometers. Other techniques like Direct Laser Writing (DLW), a method that exploits the nonlinear absorption of ultrafast laser pulses in a photosensitive material called two-photons polymerization, can also be used to construct highly intricate 3D features as small as 100 nanometers^16–18^, but their use as PDMS molds in particular remains limited.

In this paper we propose to investigate and compare the fabrication of microfluidic molds and sliding elements using both SLE and DLW methods. Such techniques have opened up exciting new possibilities for designing and manufacturing microfluidic devices, ushing beyond the limitations of traditional methods and enabling structures that would be extremely difficult, if not impossible, to produce with conventional lithography techniques. Last, we demonstrate that basic sliding elements can be created by SLE and functionalized into embedded microsensors suitable for various monitoring. As a proof of concept, we integrate microelectrodes for pH measurement directly onto the sliding element, paving the way for future *in situ*, reusable, real-time monitoring of biological sample metabolism ^19^.

## RESULTS AND DISCUSSION

### Microfluidic system development

We designed a proof-of-concept microfluidic device containing parts of highly different heights, as well as round and squared edges, with a sliding element to be inserted, to investigate the possibilities and limitations of DLW and SLE (Fig. 1a). This design was inspired by our recently published mechano-chemostat for biological samples^20^. Our microfluidic system featured inlets and outlets for culture medium flow, a compartment for a simple or functionalized sliding element, a culture chamber for introducing a biological sample via the sliding element, and a separate measurement chamber designed for sensor integration, such as pH electrodes. Notably, the electrodes were positioned away from the biological sample to avoid direct contact. The system was interconnected by microchannels that enabled continuous medium circulation. Despite its seemingly simple design, the microfluidic system incorporated channels with significant dimensional variation (see Fig 1.b), ranging from 10 µm × 10 µm square microchannels to a 480 µm × 480 µm compartment for the sliding element. Additionally, we introduced a curved geometry to prevent the formation of “dead zones” where flow and electroactive species may stagnate, potentially compromising electrochemistry measurement accuracy. This curved structure mainly allowed us, at this step, to demonstrate the fabrication capabilities. Conventional mold fabrication techniques were inadequate for such complex architectures due to their geometric limitations and susceptibility to alignment errors, in the micrometer range, which can create leaks when using sliding elements. Consequently, we adopted advanced microfabrication approaches better suited to these design requirements.

**Figure 1:**
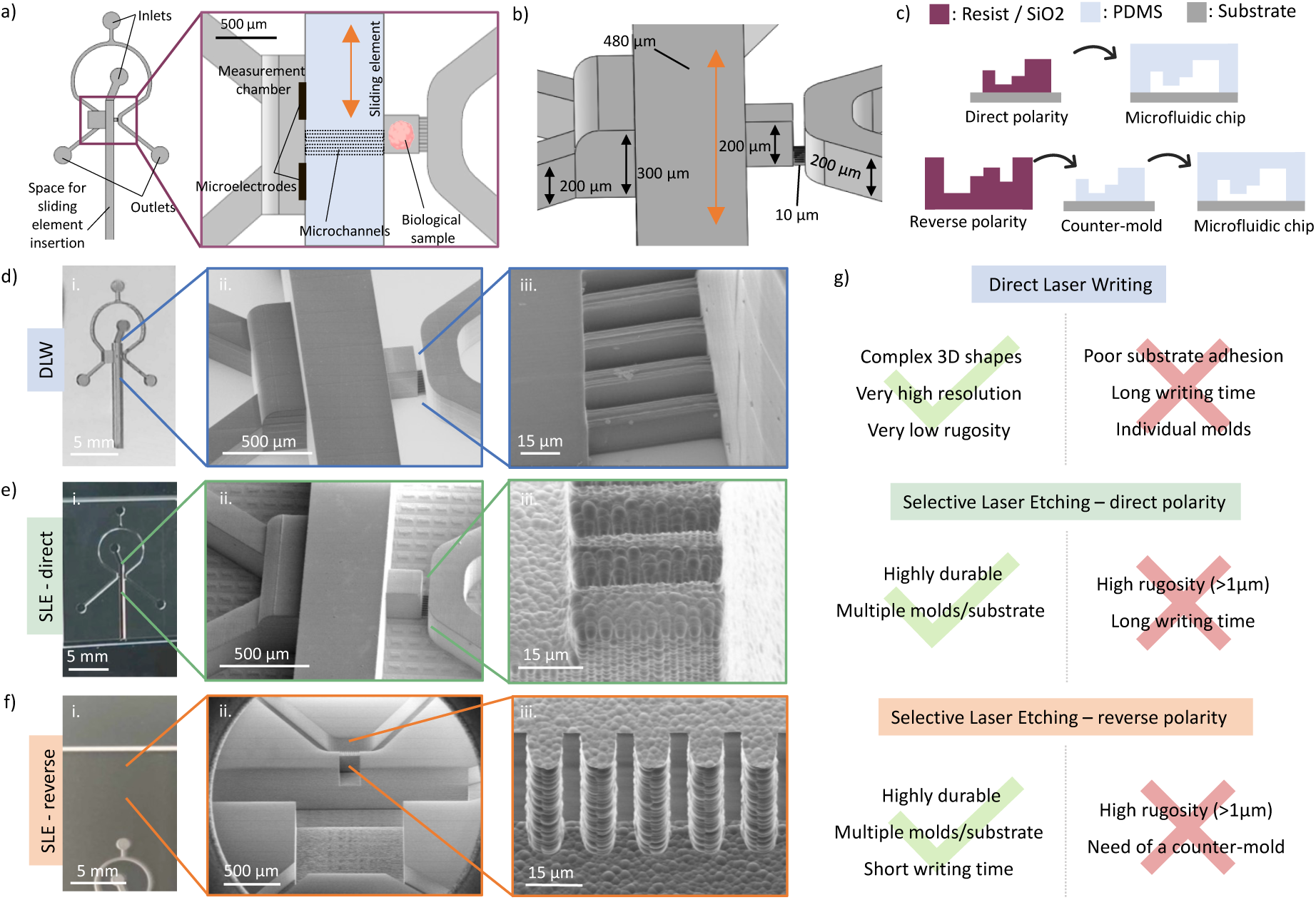
Overview of the microfluidic system design and mold fabrication developments. a) Schematic of an example microfluidic system design, comprised of a space to insert a sliding element inside the chip, with a focus on the probing area, in the case of further integration of microelectrodes onto the sliding element. b) Schematic of the microfluidic system used in this study, highlighting the different height levels required, ranging from 10µm to 480µm. The curved surface of the measurement chamber can also be observed. c) Schematic of the possible polarities regarding the mold fabrication. A direct polarity allows for fabricating microfluidic chip mold in one step, whereas the reverse polarity implies the use of a counter-mold. d) Picture of the resist mold fabricated using two-photon lithography (i), with SEM images focusing on the electrodes chamber, biological sample culture chamber (ii) and microchannels (iii). e) Picture of the positive fused silica mold fabricated using selective laser etching (i), with SEM images focusing on the electrodes chamber, biological sample culture chamber (ii) and microchannels (iii). f) Picture of the negative fused silica mold fabricated using selective laser etching (i), with SEM images focusing on the electrodes chamber, biological sample culture chamber (ii) and microchannels (iii). g) Highlights of the advantages and drawbacks of each mold fabrication method.

Fabrication steps with DLW included depositing a 500nm layer of SU-8 resist onto the surface of a silicon substrate to promote adhesion of the mold, laser-writing the direct mold structure using IP-S resist (*Nanoscribe Photonic Professional GT+*), developing the piece in SU-8 developer for 10 mins and strengthening the mold by 15 mins additional UV curing at 405nm to complete reticulation. The obtained mold showed very low rugosity (typically below 10nm), very high resolution, and easy fabrication of curved surfaces, as presented in Fig.1.d. A self-assembled monolayer of perfluorodecyltrichlorosilane (FTDS) was then grafted onto the surface of the resist mold to prevent PDMS adhesion. PDMS was cast onto the mold and cured at 65 °C for 4h. Once cured, the PDMS cast was removed from the mold and bonded to a glass substrate (see Fig.1.c), which created the channels of the microfluidic system. Yet, DLW presented some drawbacks. First, the writing time of individual molds may vary depending on the objective used (10x or 25x). In this study we decided to write the piece using the 25x objective, for optimal resolution. Consequently, the writing time was quite long (∼10 hours). In addition, the adhesion of the mold onto the substrate could be improved, as the it starts showing signs of detachment after a few PDMS castings (maximum 3 in our case), limiting its durability.

We conducted the fabrication with SLE (Nanofactory from FEMTIKA) through two different approaches: direct and reverse polarity writing. The direct polarity approach involved writing the negative space around the mold structure onto the silica piece, and performing subsequent chemical etching to reveal the direct mold, as shown in Fig.1.e. Then, the mold was silane-treated and PDMS could be casted onto the surface. Although time consuming (laser-exposing the entire negative space typically took 7h, while the chemical etching step of our large piece typically takes 5h, but can be parallelized), the mold obtained with this direct approach was highly durable and could be used for many subsequent PDMS castings (> 10 in our case). We noted a roughness of below 1 μm in our case (Fig.1.e.iii). This roughness can be decreased through a second etching^21^. We note that this roughness is not a limiting factor in our case, as it did not prevent de-molding of the PDMS, did not cause any issues during the binding of the PDMS, and did not lead to any leakage.

Additional steps were required in order to conduct the reverse polarity approach (Fig.1.f). Specifically, SLE was used to define the mold geometry within the substrate, and subsequent chemical etching selectively removed the laser-modified regions to reveal the negative mold shape (as presented in Fig.1.c). Then, the negative mold was silane-treated and PDMS was casted, to obtain a counter-mold. The counter mold was subsequently silanized and PDMS was cast again. Finally, the PDMS cast was bonded to a glass slide, creating culture chambers and channels. Advantages of this method included significantly reducing both fabrication time (etching time remains similar): laser writing the entire piece took only about one hour. However, the additional need of a counter-mold can be tedious, as counter molds often needs to be replaced due to a loss in resolution over the uses. Depending on the needs, direct polarity could thus be better suited, and not much more time-consuming.

All advantages and drawbacks of both DLW and our two approaches for SLE were summarized in Fig.1.g. Depending on the durability, size/time of fabrication, rugosity demand, which is often not a limiting factor (1μm rugosity does not limit the use of the microfluidic chip presented in Fig. 1) and resolution, both techniques offer pros and cons. We believe that SLE of direct polarity of needed microfluidic could offer more advantages than disadvantages, the difference in fabrication time being important but not human-intensive. In the following, we focused on SLE as our method of choice for microfabrication.

### Fabrication of versatile glass sliding elements

We previously demonstrated the beneficial use of the sliding elements technology in microfluidic systems for a variety of applications, including: creating reconfigurable confining culture chambers, loading and retrieving biological samples or mimicking micro-pipettes aspiration while achieving great chemical and mechanical control, and increased throughput with respect to standard aspiration^22^. First-generation sliding elements were made out of two levels of dry film sheets ^20^. Consequently, such devices could suffer from misalignment between the films, in the micrometer range, which could induce leakage and / or microchannels defects. In addition, the dry film presented natural auto-fluorescence due to the photoresist, which interfered with further fluorescence imaging experiments. Hence, we decided to optimize the device structure and material by structuring them in fused silica using SLE. Fused silica is highly durable and transparent, and that should not exhibit auto-fluorescence compared with expoxy resist-based dry films. As a consequence, the optimized devices allow optical inspection of the cells as well as the possibility to use optical markers in the cell culture parallel to the microsensors.

The sliding elements fabricated here were composed of a long and thin strip with a biological sample insertion and retrieval area (as illustrated on Fig.2.a). Fabrication included laser-writing the shape of multiple sliding elements onto a fused silica substrate (Fig.2.b) by SLE. Such elements could then be inserted into microfluidic chips, as shown in Fig.2.c, to create systems tailored for various applications. For instance, spheroids could be inserted into culture chambers, as illustrated in Fig.2.d. To this aim, the sliding element was inserted and the loading area was aligned with the culture chamber. Biological samples immersed in culture media were then injected at the inlet and guided through the culture chamber thanks to the micro flux. Last, the chamber was closed by moving the sliding element upwards, and the microchannels were aligned with the culture chamber, which allowed for the media to flow in the microfluidic system.

**Figure 2:**
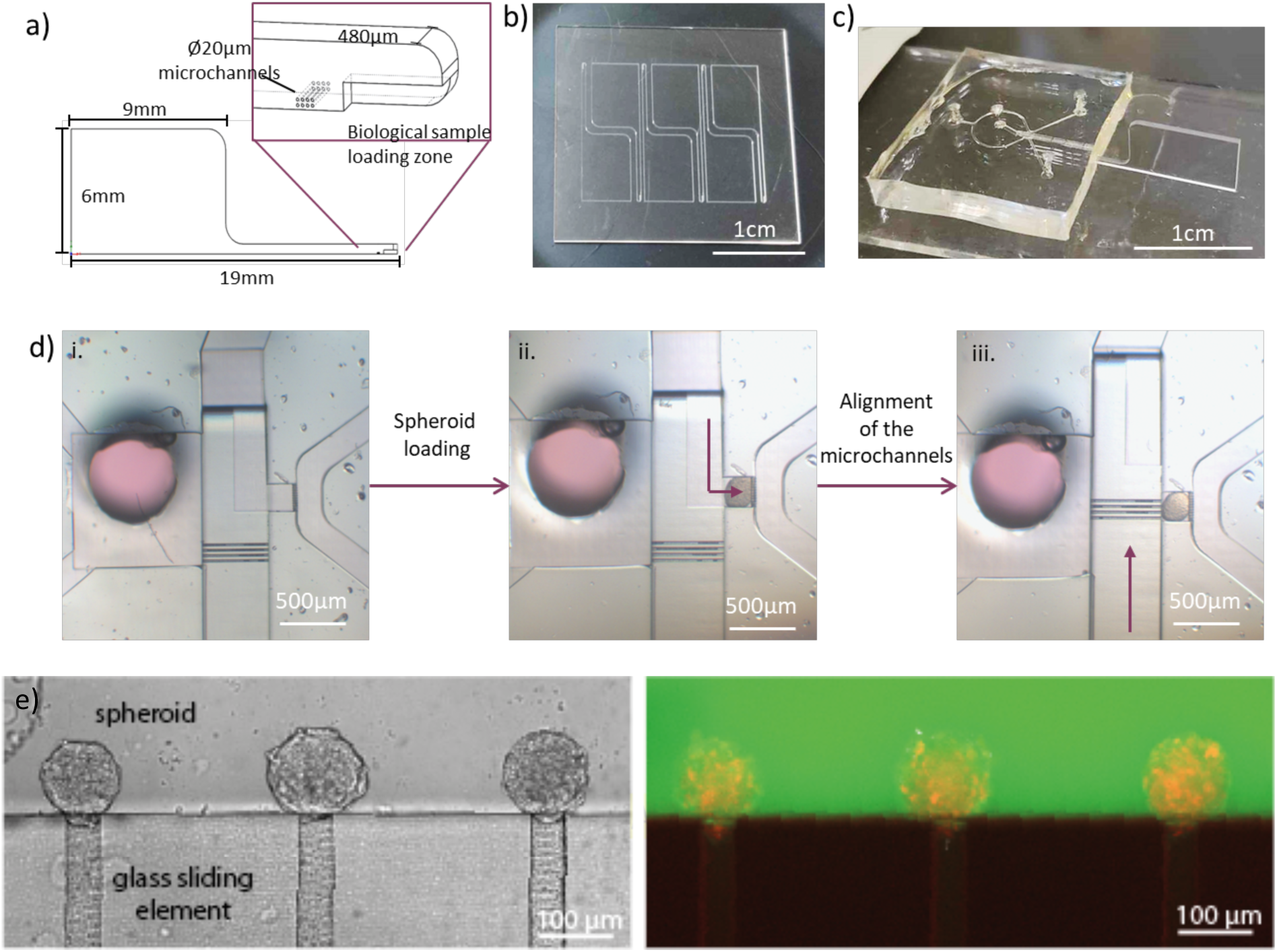
Fabrication of versatile glass sliding elements for microfluidics applications. a) Schematic of a simple glass sliding element for inserting biological samples into the microfluidic device. b) Picture of the sliding elements laser-written on a fused silica substrate. c) Picture of a glass sliding element inserted into a microfluidic device. d) Optical microscopy images showing an example of biological sample (here, a spheroid) insertion into the microfluidic device. First, the loading zone of the sliding element is aligned with the culture chamber (i). Then, flow is applied to insert the biological sample into the chamber (ii). Last, the microchannels of the sliding element are aligned with the culture chamber, closing the chamber and allowing for culture media to flow in and out of the chamber. e) Images of glass micropipette inserted in a PDMS chip. Spheroids are made of cells with a red-fluorescent tag on the nucleus, and a FITC dye is added to the culture medium. We can see that the sliding element is not autofluorescent in both the green and the red channels.

Additionally, we revisited the fabrication of the sliding elements used for micropipette aspiration on chip using SLE^22^. Similarly, we showed that we could fabricate with similar solution these sliding elements (Fig. 2e). Moreover, we observed no auto-fluoresence in both red and green channels typically used in biology (Fig. 2e right). These results together demonstrated that SLE could be efficiently used to fabricate complicated 3D shapes, from mold to sliding elements.

### Integration of micro-electrodes onto glass sliding elements for pH measurements

We proposed to convert basic sliding elements into embedded microsensors suitable for many applications, and in particular reconfigurable electrochemistry-on-chip, to perform multi-physics measurements on similar biological system. We chose as a demonstrator potential cell monitoring with the development of a pH sensor, which can be used to probe metabolic changes in cells. As a proof of concept, we integrated microelectrodes for pH measurement directly onto the sliding element, paving the way for future in situ, reusable, real-time monitoring of biological sample metabolism ^19^.

Fig.3.a illustrates the design of the device. The microelectrodes were located at the end of the tip, near the biological sample loading area, culture chamber and connected microchannels for in situ pH measurements, and the associated contact pads were placed onto the large fused-silica area. The sensor comprised a working electrode (WE) (Ø100µm) and a pseudo-reference electrode (Ø220µm) for measuring the pH in a 2-electrodes configuration, which were located above the microchannels connected to the culture chamber (Fig.3.c). This fully embedded system was fabricated using a combination of conventional lithography techniques and innovative approaches, as illustrated in Fig.3.b. and described in the materials and methods section. First, the sliding elements shapes were laser-written on a fused silica substrate. Then, the electrodes were structured through a conventional photolithography process, metal deposition and lift-off. The surface of the device was then insulated and only the recording area of the electrodes were opened using reactive ion etching. The elements then needed to be separated, yet, the liberation of individual sliding elements required a harsh chemical etching step, which would damage the exposed metallic surfaces. To overcome this limitation, a second layer of 1µm of insulator was deposited, acting as a protective layer during the chemical separation. Last, the working part of the electrodes was re-opened, the remaining sacrificial Si3N4 was removed, via a second reactive ion etching process, and the unitary sliding elements were integrated vertically into printed circuit boards (PCB) using micro welding, as shown in Fig.3.d. The functionalized sliding elements can then be insterted inside the microfluidic chip designed in Fig. 1 (Fig. 3d).

**Figure 3:**
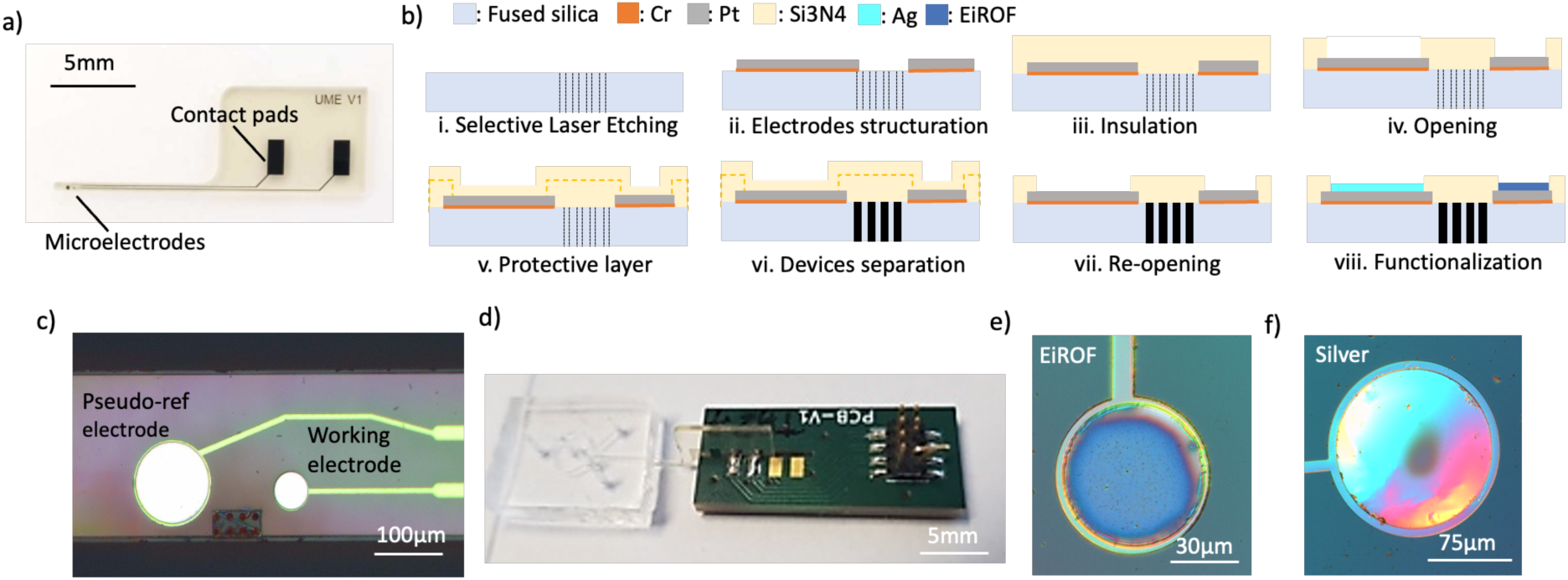
Integration of micro-electrodes onto the sliding elements. a) Picture of single sliding elements with integrated microelectrodes. b) Schematic of the microfabrication process main steps. First, the sliding elements shape are laser-written on a fused silica piece (i). Then, the electrodes are structured through a conventional photolithography process, metal deposition and lift-off (ii). The surface of the device is then insulated, (iii) and only the recording area of the electrodes are opened using reactive ion etching (iv). Next, a second layer of insulator is deposited (v), acting as a protective layer for the metallic surfaces during the chemical etching used to separate the unitary sliding elements from the fused silica substrate (vi). Finally, the working part of the electrodes are re-opened via reactive ion etching (vii) and the sliding element is ready for packaging and backend processes, which include electrochemical deposition to functionalize the electrodes (viii). For further details regarding the microfabrication process, please refer to the materials and methods section. c) Optical microscope image of the tip of the sliding element, highlighting the placement of microelectrodes around the microchannels d) Picture of a sliding element welded onto a custom-made PCB. Inset: functionalized sliding element inserted into a microfluidic chip. e) Optical microscope image of the working electrode functionalized with an EIROF layer. f) Optical microscope image of the pseudo-reference electrode functionalized with an Ag layer.

Further backend processes are required. In fact, the WE was functionalized with an Iridium Oxide layer, and the pseudo-reference electrode was covered with silver^23,24^ to perform further pH measurements. Iridium oxide is a metallic material with a wide range of oxidation states^25^, making it highly responsive to its chemical environment while maintaining excellent stability in both acidic and alkaline conditions^26^. Its diverse applications include neural interfacing, pH sensing^27,28^, catalysis^26^, cell interfacing^29,30^ or contaminant detection^31^. In this work, we focused on the electrodeposition method to modify the WE with electrodeposited iridium oxide films (EIROFs) for pH measurements applications. This technique offered several advantages when compared to other iridium deposition processes: it was accessible, operated at room temperature, did not require costly iridium substrates, and it enabled localized deposition^32,33^. According to Baur et al. ^34^, optimal EIROF coverage could be identified visually by the presence of a bright blue film. We optimized the deposition conditions (see the Materials and Methods section) to meet this criterion, as shown in Fig.3.e. Regarding the electrodeposition of the silver layer, we optimized the deposition conditions (see Materials and Methods section), to achieve a dome-shaped, shiny and reflective layer, as illustrated in Fig.3.f. Together these microfabrication steps demonstrated the first realization of an electrode on a sliding element, opening the way to reconfigurable electrochemistry-on-chip.

### Performances of the electronically-active sliding elements for pH sensing

To characterize the sensitivity of our sensor, we measured the open circuit potential (OCP) between the working electrode and a reference electrode across the range of pH values 8-6 with 0.2 pH decrements. We specifically examined how the choice of reference electrode influenced the measurements, as part of our effort to assess the feasibility of a fully embedded sensing system. For this purpose, we compared the sensor’s potential response using three different reference configurations: a commercial leakless Ag/AgCl electrode (*LF-1.6-48, Innovative Instruments, Inc*.), a bare silver wire (Ø500 µm), and the integrated electroplated Ag pseudo-reference electrode. Measurements were conducted in DPBS-based solutions, with pH adjusted by incremental additions of 0.1M NaOH or 0.1M HCl. To evaluate performance in conditions closer to biological environments, we also repeated the measurements in unbuffered DMEM-based solutions, similarly pH-adjusted, anticipating future applications in complex culture media. DMEM is the basic culture medium used in cell biology. We focused on alkaline-to-acid sequences, to mimic the cells production of lactate, by decrementing the pH value of 0.2 every 2 minutes at 25°C. Fig.4.a and Fig.4.b illustrate the OCP using the external leakless Ag/AgCl reference electrode, in DPBS- and DMEM-based solutions across the pH range 8-7. In DPBS-based solutions, the measured potential remained stable at each pH value, whereas in unbuffered DMEM, a slight drift was observed, likely due to more complex chemical interactions and the presence of electroactive species in the medium. Fig.4.c summarizes the sensitivity obtained for each condition (reference used and solution), and results appeared to be dependent on the reference electrode rather than the solution. For instance, when using the leakless Ag/AgCl electrode, we obtained sensitivities of 65 ± 4mV/pH (n=6) in DPBS and 63 ± 4mV/pH (n=3) in DMEM. With the Ag wire, we obtained 86 ± 12mV/pH (n=6) in DPBS and 85 ±11mV/pH (n=7) in DMEM. Last, with the integrated pseudo-reference electrode we obtained 66 ± 10mV/pH (n=3) in DPBS and 56 ± 5mV/pH (n=3) in DMEM.

**Figure 4:**
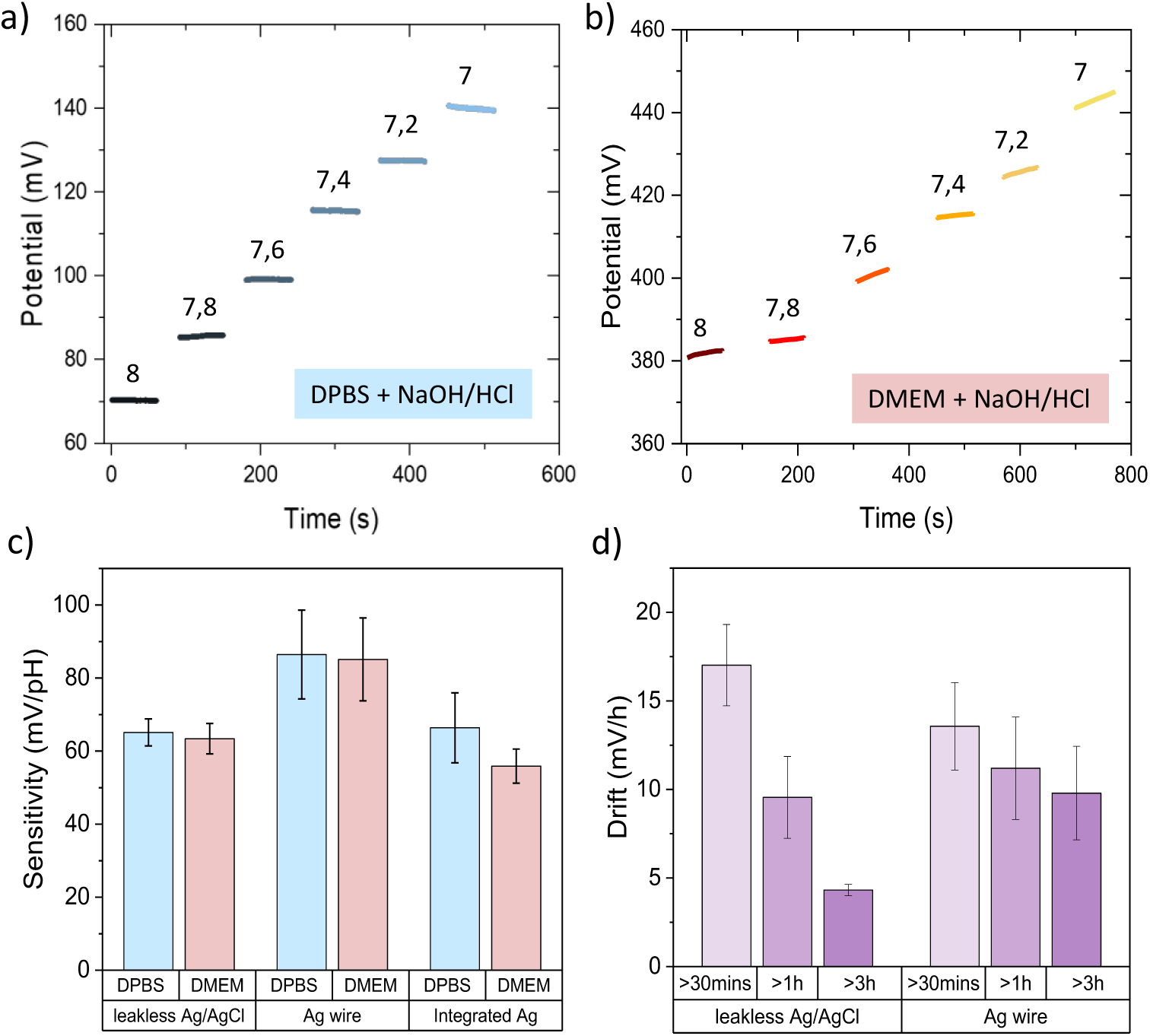
Performances of micro-electrodes for pH sensing. a) Graph illustrating the potential response of the WE measured in an OCP configuration, using an external leakless Ag/AgCl reference electrode. The potential steps correspond to different pH values in DPBS-based solutions. b) Graph illustrating the potential response of the WE measured in an OCP configuration, using an external leakless Ag/AgCl reference electrode. The potential steps correspond to different pH values in DMEM-based solutions. c) Bar chart showing the sensitivity of the WE, in mV/pH, depending on solution and reference electrode used. d) Bar chart indicating the drift of the measured potential over time in DMEM, when using external reference electrodes.

To further evaluate the performance of our electrodes, we investigated signal drift over time under physiological conditions, anticipating their future use in complex culture media. Our goal was to assess the measurement stability, with particular attention to the first three hours, an important window during which significant pH fluctuations were expected due to the very small media volumes typically used in microfluidic chips. For this purpose, the WE was immersed in complete culture medium (pH 8; full composition detailed in *Materials and Methods*) for up to four hours. After an initial 30-minute stabilization period, we quantified the potential drift over three intervals: from 30 minutes to 1 hour, after 1 hour, and after 3 hours of immersion, as shown in Fig.4.d. The following results were obtained: When using the external leakless Ag/AgCl electrode, drifts were equal to 17 ± 2mV/h (n=4), 9 ± 2mV/h (n=3) and 4.3 ± 0.3mV/h (n=4) for 1 hour, after 1 hour, and after 3 hours of immersion, respectively. The measurement drift seemed to stabilize over the first 3h to reach a value of 4mV/h. Note that the maximum drift remained about 10% of the measurement over short timescale, to drop to a few percents at longer timescales. On the other hand, when measuring the OCP between the WE and a simple Ag wire, drifts were equal to 13 ± 2.mV/h (n=5), 11 ± 3mV/h (n=5) and 10 ± 3mV/h (n=4) for 1 hour, after 1 hour, and after 3 hours of immersion, respectively. The stabilization of the drift appears steady yet much slower when compared to commercial reference electrodes.

### Conclusion and perspectives

In this paper, we presented the use of DLW and SLE as alternatives to create both high resolution molds and sliding elements for reconfigurable microfluidic devices. While DLW provided the higher resolution and lower rugosity, it took longer to create molds and the mold detached after some use. Conversely, SLE, in particular if performed on the reverse polarity, was quick and highly durable. Both DLW and SLE allowed for the fabrication of complex 3D molds, with curved surfaces, and different heights.

Using SLE, we demonstrated that we could also fabricate sliding elements, to be inserted inside a microfluidic device. Moreover, we demonstrated that SLE could be coupled to a classical micro-electronic process and electrodeposition to create functionalized sliding elements for reconfigurable electrochemistry-on-chip: the sliding elements can be written on a glass wafer first, then classical planar electro-fabrication can be performed before etching the sliding elements. A great attention had to be paid on the compatibility and sequence of the different fabrication steps, especially with regards to the KOH used at the end of the process to detach the sliding elements from the wafer. Yet, we demonstrated that this limitation can be overcome by the addition of a protective layer, to prevent damaging the surfaces during this last etching step.

Our sensors overall exhibited a very high sensitivity compared to the literature, as other IrOx-based pH sensor usually exhibit responses ranging from near-Nernstian (around 50mV/pH) to super-Nernstian (above 59mV/pH and up to 69mV/pH) at 25°C^27,35–39^. Yet, very few pH measurements studies had been conducted using micro electrodes and were usually performed with electrodes of greater area ^40^, hence, results regarding microelectrodes were quite novel. Nonetheless, studies performed on microelectrodes (from Ø10µm tips to 50×100µm rectangles) also exhibited a super-Nernstian sensitivity of 70-80mV/pH^41^ and 74.2mV/pH ^42^, respectively.

Overall, the drift values we obtained were higher than those reported in literature, however typically measured on electrodes of greater area. Nonetheless, Nguyen et al. performed studies on microelectrodes (50×100µm) and obtained a drift of 72mV/h at pH 7.78 which is higher than what our microsensor exhibits. Overall, studies that have explored pH measurements using Iridium-coated microelectrodes, usually rely on larger-area electrodes and bulky reference systems such as calomel electrodes. As a result, our findings represent a novel contribution to the field, particularly in the context of miniaturized, embedded sensing platforms.

To conclude, we showed in the paper that SLE is a very good candidate to create novel 3D-molds and sliding elements and that these ones can be functionalized to create electrodes. The combination of these two leads to reconfigurable microfluidic and electrochemistry-on-chip: while the sliding element can bring novel functions that can be coupled to deformable elements inside a microfluidic device to mechanically and chemically control the biological sample, the inserted electrode can be used to perform additional measurements, paving the way to multi-physics analyses of biological samples.

## MATERIALS AND METHODS

### Direct laser writing with Nanoscribe

Our direct laser writing process operates on two-photons exposure. To manage this technique, we use Photonic Profesional GT+ equipment and IP-S resin (Nanoscribe). The main process parameters are fixed with Describe software (Nanoscribe) and an STL file of the mold. The total height of the mold is 480 µm. It is sliced with 240 layers. The mold’s length and width are 13.6 mm and 6.5 mm, respectively. It is assembled from 697 blocks. Each block is scanned by a pulsed infrared laser with a 25x optical lens. The block connection positions and block overlaps are optimized to increase the mold quality. The main options for configuring block writing are:

– Hatch spacing: 0.5 µm
– Writing mode: shell and scaffold
– Interface detection: automatic for critical blocks with small patterns
– Writing speed: 100 mm/s

### Selective laser etching with Femtika Laser Nanofactory

We used a Femtika Laser Nanofactory system (Femtika.Ltd, Lithuania), equipped with an ultrafast Yb:KGW femtosecond laser (1030 nm central wavelength, 700 fs pulse duration, 610 kHz repetition rate), to expose glass substrates.

High-precision XYZ translation stage combined with galvanometric scanners enables to focus the laser beam into the volume of the glass substrate at high precision. We used a 20× microscope objective with a numerical aperture (NA) of 0.45, which produced a point spread function of about 3 µm in diameter in the XY plane and 14 µm along the Z-axis. The writing process was performed in an ambient air condition, on a vibration-isolated stage to maintain alignment stability.

After the laser-induced modification process, the substrate was plunged into a potassium hydroxide (KOH) solution (6M) at 90 °C for 5 hours. Following this step, the substrate cooled down in the oven until it reached 50 °C. The substrate was cleaned with deionized (DI) water and left to dry in ambient conditions.

### Microfabrication process of the electronically-active sliding element

First, the sliding elements shape and channels are laser-written on a fused silica piece (Nanofactory Femtika), as illustrated in Fig.2.b. The fused silica substrate is subsequently cleaned using buffered oxide etch (*BOE 7:1, MicroChemicals GmbH*), to remove any silica dust or particles resulting of a small ablation during the writing when the laser beam is at the interface air/glass. Then, the electrodes shape, contact pads and access lines are structured using photolithography (*MABA6Gen4, Suss Microtec*), subsequent Cr (50nm) and Pt (200nm) physical vapor deposition (*Eva600, Alliance Concept*) and lift-off of the resist in acetone. Then an insulation layer of SI_3_N_4_ (1µm) is deposited using plasma-enhanced chemical vapor deposition (*PECVD100, ApSy*). thickness of the insulation layer allows for both electronical insulation for the liquid media and mechanical shield as the sliding-element will further be inserted into and retrieved from the microfluidic system. Next, the recording area of the electrodes are opened using reactive ion etching (*SI500-DRIE, Sentech*). The next step consists in separating the sliding elements from the fused silica substrates using a KOH-based chemical etchant. Prior to this step the sliding elements are covered in an additional 1µm of Si_3_N_4_, acting as a sacrificial layer, to protect the metallic surfaces from the etchant. Last, the residual Si_3_N_4_ is removed from the recording area of both electrodes through the same plasma etching process, and unitary sliding elements are ready for packaging. To this aim, each device is inserted in the slit of our custom-designed PCB and contact pads are welded, as shown on Fig.3.d.

Focused-ion beam (FIB) milling and scanning electron microscopy (SEM) were performed with a HeliosNanoLab 600i Dual Beam (*FEI*) for characterization of the surfaces.

### Functionalization of the working electrode

To perform EIROF deposition we used the solution developed by Marzouk et al.^43^, following the pioneering work of Yamanaka et al.^44^. The EIROF depositions are performed in a conventional 3-electrode setup, with a Pt wire as the counter electrode, an external commercial leakless Ag/AgCl electrode as the reference electrode (*LF-1.6-48, Innovative Instruments, Inc*.) and the integrated Pt working electrode. We use the potentiostatic route, with an applied potential of 0.6V until the desired charge of 25mC/cm² is consumed.

### Functionalization of the pseudo reference electrode

To perform silver deposition, we used a commercial electrolyte solution (*Silver electrolyte, TIFOO.)* and silver has been deposited using the galvanostatic route in a 3-electrode configuration as described above. We applied a current of 1µA for 15mins.

### Solutions

pH measurements were conducted either in DPBS-based (ThermoFisher) solutions or in unbuffered DMEM-based solutions (ThermoFisher), and were pH adjusted by adding small amounts of 0.1M NaOH, or 0.1M HCl. The complete culture media used for drift measuring is composed of unbuffered DMEM supplement with fetal bovine serum (FBS), penicillin / streptomycin, glutamax and pyruvate.

## ACKNOWLEDGMENTS

We thank Jérôme Launay, Pierre Temple-Boyer, Bastien Venzac, Moetassem Meksassi and Juliette Lignières for their help and discussion in the elaboration of this project. This work was supported by LAAS-CNRS micro and nanotechnologies platform, a member of the Renatech French national network, by Nanofutur project, funded by the ANR (grant agreement 21-ESRE0012), the Inserm Plan Cancer MechaEvo, and the ERC Starting Grant *UnderPressure* (grant agreement number 101039998). Views and opinions expressed are, however, those of the author only and do not necessarily reflect those of the European Union or the European Research Council. Neither the European Union nor the granting authority can be held responsible for them.

